# A Neural Network for High-Precise and Well-Interpretable Electrocardiogram Classification

**DOI:** 10.1101/2024.01.03.573822

**Authors:** Xiaoqiang Liu, Liang He, Jiadong Yan, Yisen Huang, Yubin Wang, Chanchan Lin, Yingxuan Huang, Xiaobo Liu

## Abstract

Manual heart disease diagnosis with the electrocardiogram (ECG) is intractable due to the intertwined signal features and lengthy diagnosis procedure, especially for the 24-hour dynamic ECG signals. Consequently, even experienced cardiologists may face difficulty in producing all accurate ECG reports. In recent years, neural network-based automatic ECG diagnosis methods have exhibited promising performance, suggesting a potential alternative to the labor-intensive examination conducted by cardiologists. However, many existing approaches failed to adequately consider the temporal and channel dimensions when assembling features and ignored interpretability. And clinical theory underscores the necessity of prolonged signal observations for diagnosing certain ECG conditions such as tachycardia. Moreover, specific heart diseases manifest primarily through distinct ECG leads represented as channels. In response to these challenges, this paper introduces a novel neural network architecture for ECG classification (diagnosis). The proposed model incorporates Lead Fusing blocks, transformer-XL encoder-based Encoder modules, and hierarchical temporal attentions. Importantly, this classifier operates directly on raw ECG time-series signals rather than cardiac cycles. Signal integration begins with the Lead Fusing blocks, followed by the Encoder modules and hierarchical temporal attentions, enabling the extraction of long-dependent features. Furthermore, we argue that existing convolution-based methods compromise interpretability, while our proposed neural network offers improved clarity in this regard. Experimental evaluation on a comprehensive public dataset confirms the superiority of our classifier over state-of-the-art methods. Moreover, visualizations reveal the enhanced interpretability provided by our approach.

**Highlights:**

1. Our model extracts long-dependent features of ECG signals based on the Transformer-XL encoder.
2. The proposed network offers the improved interpretability.
3. Our classifier achieves superior performance over other state-of-the-art methods.

## I. Introduction

EART diseases, such as tachycardia, atrial fibrillation, and atrial premature beats, pose significant life-threatening risks. Diagnosis of these conditions relies on electrocardiogram (ECG) signals recorded by specialized monitoring devices. However, the interpretation of ECG signals for diagnoses, particularly conditions like atrial fibrillation, is a labor-intensive and intricate process. This complexity arises from the inherent noise present in ECG signals and the fact that features of heart diseases are often subtle and entangled within the ECG data. Analogous to the manual analysis of various time-series data, ECG signal interpretation necessitates a meticulous examination of diverse waveforms. This process lacks the intuitive clarity afforded by the analysis of two-dimensional images. Consequently, there is a pressing need for accurate ECG signal classification methods tailored to specific heart diseases.

In recent times, increased attention from scholars has been directed towards methods for automating heart disease diagnosis. Notably, automatic diagnosis (classification) methods [1]– [5] have demonstrated significant performance advancements leveraging machine learning techniques. These methodologies hold potential to either supplement or potentially assume certain responsibilities of cardiologists, particularly in scenarios like the continuous monitoring inherent in dynamic ECG, often necessitating 24-hour observation. Initially, automatic ECG diagnosis methods involved a phase of feature extraction through statistical summarizations [6], [7], followed by a classification stage employing classifiers such as the support vector machine [8] or the multi-layer neural network [7]). While manually engineered features may offer reasonable insights into signal analysis, the paradigm of automated feature extraction has shown to be more adept at ensuring accurate classification [1], [9].

Neural networks have demonstrated significant superiority over traditional machine learning methods in various domains, including image recognition [10], speech recognition [11], natural language processing [12], [13], medical image analysis [14]–[16], and point cloud data analysis [17], [18]. Given the global repository of over 300 million recorded ECGs [19], these vast datasets provide an ideal foundation for training large-scale neural networks. End-to-end neural networks [1], [3], [20]–[22] have been effectively applied to ECG signal classification, seamlessly integrating feature extraction and classification within a unified framework. Hannun et al. [9] introduced a straightforward neural network for ECG classification, achieving performances comparable to human cardiologists. Several other neural network approaches [1], [3], [4], [23] have been proposed, each offering unique perspectives, such as data augmentation, multi-label regularization, and robust feature extraction. Additionally, attention-based methods [24] have proven valuable in feature extraction, enhancing interpretability.

Nonetheless, two key considerations must be addressed in designing an effective automatic classifier: high precision and interpretability. Prior efforts have often neglected the modeling of long-term dependencies in ECG signals. This is particularly critical when dealing with extended time-series data, such as the continuous 24-hour signals recorded by dynamic ECG monitoring. Neglecting long-dependent features can hinder the accurate diagnosis of heart diseases. For instance, tachycardia, defined as a state with an average heart rate exceeding 100 beats per minute, requires extended observation for diagnosis. Clinical practice also demands the scrutiny of several cardiac cycles to diagnose atrioventricular block. Furthermore, mainstream methods have several cardiac cycles to diagnose the atrioventricular block. Additionally, the known mainstream methods extracted features by convolutions that are essentially weighted moving average operations. Summary features were extracted through the average, but the exact location information is vanished.

In this paper, we present a novel neural network framework designed for ECG signal classification, incorporating both local feature extraction and long-dependence capture. After initial pre-processing steps, such as denoising, the ECG signals are input into a series of Lead Fusing blocks. These blocks consist of multiple 1D convolutions with ReLU activation [25] and incorporate the Squeeze-and-Excitation Attention mechanism [26]. Their purpose is to extract time-aligned features and fuse the ECG signals in the channel dimension.

Following this initial feature extraction stage, the processed data is then forwarded through two stacked Encoder modules. Each Encoder module employs a sliding window approach to partition the learned time-aligned features into smaller local and real-time segments in the length dimension. The Transformer-XL encoder, a variant of the Transformer architecture [13], is then applied to these segments to extract non-local features. Notably, the Transformer-XL encoder is capable of capturing features from both the current and previous receptive segments.

Certain diseases can only be accurately diagnosed by closely observing specific cardiac cycles. To address this challenge, we introduce a hierarchical temporal attention module that plays a crucial role in discerning critical non-local features and acquiring a comprehensive understanding of global features.

Subsequently, these global features are directed into a fully connected layer for classification, following a methodology similar to prior research [1], [9].

Experimental results demonstrate that our proposed neural classifier surpasses the performance of previous state-of-the-art methods on a public ECG dataset comprising a large collection of multi-lead ECG signals. Furthermore, compared to previous convolution-based approaches, our framework employs fewer convolutions, ultimately enhancing interpretability, which is corroborated by GradCAM [27] visualization techniques.

This work presents several significant contributions:

- We introduce a novel neural classifier for ECG signal classification (diagnosis), which leverages the Transformer-XL encoder to extract non-local features and model long-term dependencies effectively.
- We incorporate a hierarchical temporal attention module to consolidate the features derived from the Encoder modules and predict global features.
- Experimental evaluations conducted on the public Tianchi ECG dataset confirm the superior performance of our proposed neural classification method in comparison to previous state-of-the-art approaches.
- Our approach enhances interpretability by minimizing the use of convolutions on ECG data, resulting in clearer and more intuitive insights.

## II. Preliminaries and Related Work

### A. Backgrounds on ECG signals

ECG signals are non-invasively recorded through metal leads, providing representations of heartbeats [28], [29]. Various ECG monitoring systems, including the Holter, event, patch, and implantable monitors, are employed in different clinical scenarios. ECG recordings typically consist of multiple signals from different leads, akin to the multiple color channels captured by cameras or human eyes in natural images. Following the methodology of previous studies [1], [9], ECG signals from different leads are treated as distinct data channels. As illustrated in Fig. 1, we showcase an example of normal ECGs recorded via 12 leads from the ECG library^1^. Each segment of ECG signals encompasses multiple cardiac cycles, each representing a complete heartbeat. However, some diseases can only be accurately diagnosed by analyzing more than one cardiac cycle, such as atrioventricular block and tachycardia. Therefore, the diagnosis of heart disease necessitates an examination of a relatively extended period rather than just one or a few cardiac cycles. Given this unique data structure, our focus for accurate classification centers on two key aspects: i) The fusion methods to analyze signals from different leads; ii) Long-dependence modeling to capture non-local and global features.

**Fig. 1.**
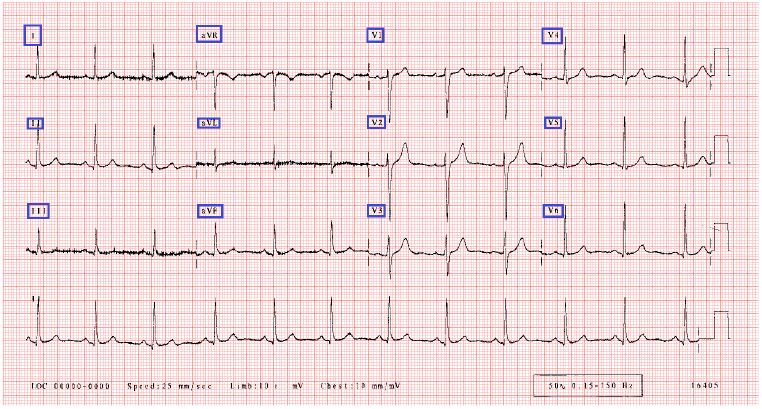
An example of normal ECG signals. The sign marked in blue rectangles indicate which which leads record the signal.

### B. Transformers

Initially, Transformers [12], [30] exhibited their efficiency in natural language processing [31], [32], before finding application in computer vision [33], [34]. In earlier research, strategies for processing time-series data were predominantly characterized by original and variant methods of recurrent neural networks [35]–[37]. In the current landscape, transformer-based approaches [12], [13] have made significant contributions by utilizing self-attention mechanisms [38] to capture data relationships.

Nonetheless, the original transformers introduced substantial computational complexities when dealing with lengthy sequences. Subsequent efforts aimed at refining these models with improved computational efficiency. The reformer [39] employed a hashing method to identify critical features before self-attention computations. Linformer [40] utilized low-rank approximation to reformulate attention weights, effectively replacing the *O*(*n*^2^) operations of the original transformer [12] with an *O*(*n*) operation. Similarly, the Performer [41] introduced a concept using orthogonal random features to eliminate the need for storing and computing attention weights. Transformer-XL [13] drew inspiration from recurrent neural networks by establishing connections between adjacent local segments. Specifically, segment-based recurrent methods, such as the Transformer-XL, demonstrated remarkable flexibility in managing long sequences.

These advances in sequential data analysis led to the introduction of some transformer-based methods [42], [43] for ECG classification, resulting in commendable performance. However, it is worth noting that these methods did not explicitly model long-term dependencies, potentially omitting critical features in the process.

### C. Neural networks for ECG classification

Numerous neural network approaches [1]–[3], [20]–[22] have been proposed for ECG classification. Rajendra et al. [20] introduced a nine-layer convolutional network designed to recognize five common heart diseases. Kiranyaz et al. [22] presented a real-time, patient-specific ECG classification system employing 1D convolutional neural networks. In [21], a more complex method was introduced, which included an R-peak detection algorithm, feature extraction through convolutional neural networks, and the application of handcrafted rules. In [4], neural networks with residual connections were employed, resulting in distinguishable learned representations for arrhythmia classification. An innovative variant of Mixup called Flow-Mixup [1] focused on correlative feature decoupling and data regularization, promoting multi-label ECG signal classification.

Generative adversarial networks (GANs) [3], [44] were utilized for data augmentation, enhancing ECG classification. Literature [9] conducted a comparison between automatic diagnosis systems based on neural networks and cardiologists, demonstrating that convolutional neural networks could achieve performance levels similar to cardiologists. Beyond convolution-based networks, Recurrent Neural Networks (RNNs) [45]–[47] proved valuable in ECG classification, highlighting the significance of sequential features. Recently, variants of the transformer model [12], [40], [41] have made substantial strides in processing sequential data and effectively classifying ECG signals [42], [43] while also incorporating handcrafted features. In [48], a comparison between a hand-crafted feature-based classifier and a neural classifier revealed that neural networks outperformed in ECG classification.

## III. Method

### A. The Review of the Transformer-XL

The self-attention module in the original transformer [12] was limited to processing data with a predefined length, resulting in high computational complexity when dealing with lengthy input data. However, Transformer-XL [13] addressed this limitation by incorporating segment-level recurrence into the attention mechanism and introducing relative position encoding to replace absolute position encoding. In Transformer-XL, the self-attention is defined as:

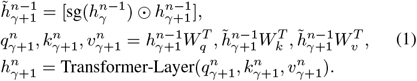

where 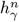 denotes the *γ*-th segment within the hidden states of the *n*-th layer, sg(·) indicates the stop-gradient operation, and [*h* ⊙ *h*] means the concatenation along the length dimension. The Transformer-Layer(consolidate the features derived from the Encoder modules and predict) stands for the self-attention computing in the original transformer. This design allows the previous segment to influence the current attention computation during forward propagation, but it does not affect backward propagation. This approach enables Transformer-XL to effectively learn long-term dependencies in sequential data.

### B. Our proposed Neural Classifier

In our work, we introduce a novel framework for heart disease classification, comprising Lead Fusing blocks, Encoder modules with Transformer-XL encoders, and hierarchical temporal attention modules, as illustrated in Fig. 2.The Lead Fusing blocks fuse the features along the channel dimension. In addition, the Encoder and temporal attention modules extract the features in the temporal dimension.

**Fig. 2.**
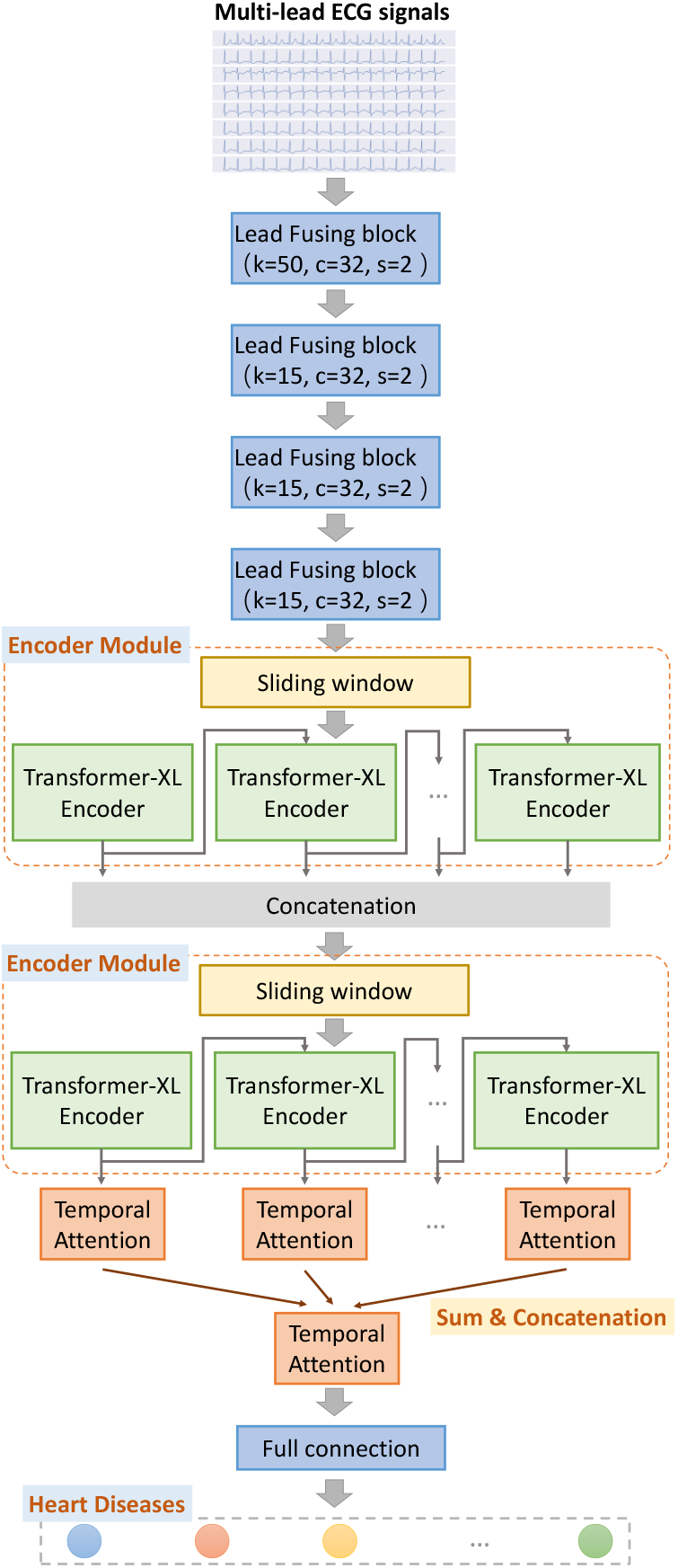
An illustration of our proposed neural classifier for heart disease diagnosis. An Encoder module is built based on the transformer-XL encoder, and the parameters of the temporal attentions are shared in the same layers. The input ECG signal is recorded by multiple leads, and the signal from each lead is treated as one input channel in our method. The parameters of the Lead Fusing Block in reported in the parentheses: “k”, “s”, and “c” indicate the “kernel size”, “stride”, and the “channel number” of the convolution in the main path (but not the convolution in the shortcut path).

#### 1) Lead Fusing Block

Following data pre-processing, the ECG signals *x* ∈ ℝ^*l×c*^ (*l* and *c* are the length and channel number) are forwarded through several Lead Fusing blocks for channel fusion. Prior approaches often processed ECG signals using 1D convolutions [1], [9], overlooking the unique relationships among different leads. While 1D convolutions can compute a weighted average over the signal leads, the use of fixed weights is impractical. Specifically, signals recorded by various leads capture the same cardiac activities from different perspectives. Each signal conveys specific information, which can be of vital importance in certain cases [49]. Moreover, the significance of these signals can vary under different circumstances.

To alleviate this concern, we introduce an innovative Lead Fusing block. In particular, following a residual learning strategy, we incorporate squeeze-and-excitation attention (SE-Attention) [26] to enhance the fusion of features from different channels, as depicted in Fig. 3. Formally, for signals (or features) 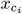 originating from lead *c*_*i*_, the output features are determined as follows:

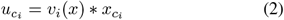

where the *v*_*i*_(*x*) squeezes the length features within the *i* channel into a scaling factor, and “*” operator denotes broadcast multiplication. This approach, in contrast to 1D convolutions, enables SE-Attention weights to dynamically assign different weights in various cases.

**Fig. 3.**
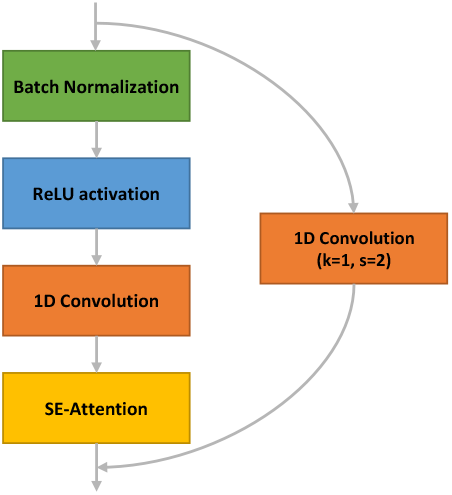
Illustrating the detailed structure of the proposed Lead Fusing block. The hyper-parameters of the Convolutions follow the hyper-parameters of the 1D convolutions in the Champion Network in the Tianchi ECG Competition. “k” and “s” indicate the kernel size and the stride step of the convolution.

We finalize this process by stacking 4 Lead Fusing blocks sequentially to process and consolidate information from multi-lead signals. The convolution kernel size is set to 50 in the first block and 15 in the subsequent ones.

#### 2) Encoder Module

The Encoder modules in our proposed framework handle the features generated from the final Lead Fusing block, as illustrated in Fig. 2. Each Encoder module is composed of a sliding window and incorporates the Transformer-XL encoder [13]. Unlike text data, ECG signals lack easily identifiable breakpoints, akin to full stops or paragraphs. In a previous transformer-based ECG classifier [42], ECG signals were processed directly by the attention mechanism, which led to elevated computational complexity. Another transformer-based approach [43] necessitated manual annotations of R peaks, and a hard division was employed in [46].

In our design, a sliding window is employed to real-time obtain overlapping segments for the Transformer-XL encoder. The length of the window is set 32 with step 24 (remaining an 8-grid overlapping fragment). This configuration is determined based on the average length of cardiac cycles. The segments obtained by the sliding window are subsequently processed by two-layer encoders of the Transformer-XL, following Eq. 1, as depicted in Fig. 2. After passing through the first Encoder module, the features are concatenated along the length dimension and forwarded to the second Encoder module. As mentioned in Sec. III-A, the non-local features are acquired within the Encoder module through the use of Transformer-XL encoders. Subsequently, the output features are directed into a novel hierarchical temporal attention module, which further integrates the non-local features to derive global features.

#### 3) Hierarchical Temporal Attention

In the Encoder module (Sec. III-B.2), we focus on learning the non-local features through the Encoder modules in the length dimension. However, we have not yet captured the global features. Since Transformer-XL encoders break down the self-attention computation into several steps, we require a global feature extractor to fuse the features from each step (for each segment). Inspired by spatial attention mechanisms, we introduce a hierarchical temporal attention operation to handle these non-local features and obtain higher-level semantics (global semantics). The operation in the temporal attention is defined as follows:

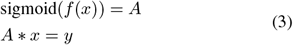

where *x* ∈ ℝ^*l×c*^ (*l* and *c* indicate the length and the number of channels, respectively), and the function *f* denotes a mapping function realized through neural operations, such as convolution with non-linear activation. The attention weight *A* in Eq. (3) shares the same dimensions as the input *x* and is applied through element-wise multiplication, denoted by the “*” operator.

In the first attention layer, the temporal attentions are parameter-shared. The length of the resulting features is reduced to a single value through averaging, and these values are concatenated before being fed into the second attention layer. This procedure can be formally defined as:

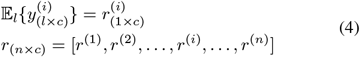

where *y*^(*i*)^ denotes the *i*-th produced features from the temporal attention, and the subscripts indicate the feature sizes. 𝔼 signifies an average operation. Therefore, the hierarchical temporal attention further consolidates the non-local features: the first attention layer processes the segment features, and the second combines these features to capture global information. Finally, the extracted features are fed into a fully connected layer to make predictions regarding ECG diseases. Similar to the previous works on multi-label classification, the model is trained using the binary cross-entropy loss function, defined as:

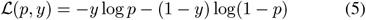

where *y* and *p* indicate the target label and predicted probability, respectively.

### C. Interpretability Analysis

In the past, convolution-based models were predominant in the ECG classification field. While moving and averaging operations can group and fuse local features to reveal smoother features, models with numerous stacked convolutions tend to obscure the precise locations of these features. To mitigate this issue, we have limited the use of convolutions and focused on feature extraction through attention mechanisms, such as SE-Attention in the Lead Fusing blocks, Self-Attention in the Encoder modules, and temporal attention.

For instance, when processing an input *x* ∈ ℝ^10^, it would require 4 stacked convolutional layers with kernel sizes of 3 and a stride of 1 to fuse the first value *x*^(1)^ and the last value *x*^(10)^. In contrast, an attention module (e.g., self-attention) may achieve the same result in just one step. Therefore, the attention-based design facilitates the extraction of long-range dependencies in ECG signals with fewer steps, preserving the precise feature locations.

## IV. Experiments

### A. Data Preparation

In this section, we will conduct experiments on the Tianchi ECG dataset^2^ to evaluate the effectiveness of our proposed method. We have selected 19,759 samples from the total 20,038 8-lead ECG records in the dataset, covering the most common 20 ECG categories, each representing a specific disease. These signals are recorded at a frequency of 500 Hertz, and each belongs to more than one category. Before feeding into the proposed networks, we first pre-processed the ECG signals, involving de-noising (by using the python package [50])) and linear scaling normalization by:

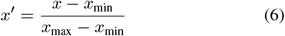

where *x*_min_ and *x*_max_ are the minimum and the maximum values of the ECG signals, respectively.

### B. Experimental Setup

We have implemented the proposed networks using Python (4) 3.8 and PyTorch 1.7 on RTX2080Ti GPUs. We set the batch size as 128 and utilize the stochastic gradient descent (SGD) optimizer [51] during the training phase. The learning rate is initially set at 0.1 and decreases 10 × when training loss does not decrease for 5 consecutive epochs. For comparison, the performances of previous works are obtained by running the reimplemented codes in their default experimental setup as specified in their original papers.

### C. Classification performances

In the comparison with state-of-the-art methods, which include the 1D ResNet-34 [52] (the Champion Network in the Tianchi ECG classification Competition^3^), ECG-Net [24], and Hannun’s model [9] (prone comparable to the cardiologists), our method exhibits superior performance across Precision, Recall, and Micro F1 indicators. These indicators are official metrics in the Tianchi ECG classification Competition. As depicted in Table I, our approach outperforms previous methodologies, including ResNet34 with Mixup [53] employed for data augmentation. Particularly noteworthy is the margin by which we surpass Hannun’s model [9] (with over a 1% increase in Precision and Micro F1, and a 0.9% boost in Recall). This suggests that our proposed method may achieve comparable or even better performance than that of cardiologists. These results also confirm that the hyperparameters utilized for the Lead Fusing blocks and the sliding windows are suitable for ECG classification.

**TABLE I.** The classification performances of the proposed methods and the current state-of-the-art work. The best performances are marked **in bold**.

| Methods | Precision | Recall | Micro F1 |
| --- | --- | --- | --- |
| Ours | **0.943** | **0.890** | **0.916** |
| Champion Network | 0.935 | 0.888 | 0.912 |
| 1D ResNet34 | 0.938 | 0.872 | 0.904 |
| 1D ResNet34+Mixup | 0.930 | 0.887 | 0.908 |
| ECG-Net | 0.911 | 0.851 | 0.880 |
| Hannun's | 0.923 | 0.881 | 0.902 |

To further compare the classification performances in each ECG category, we illustrate the ROC-AUC curves in Fig. 4. As outperforming Hannun’s [9] in most categories (except for the “Counterclockwise rotation” and “first-degree atrioventricular block” categories), our methods approach particularly excels in categories with a substantial number of samples, demonstrating its superior performance in those cases.

**Fig. 4.**
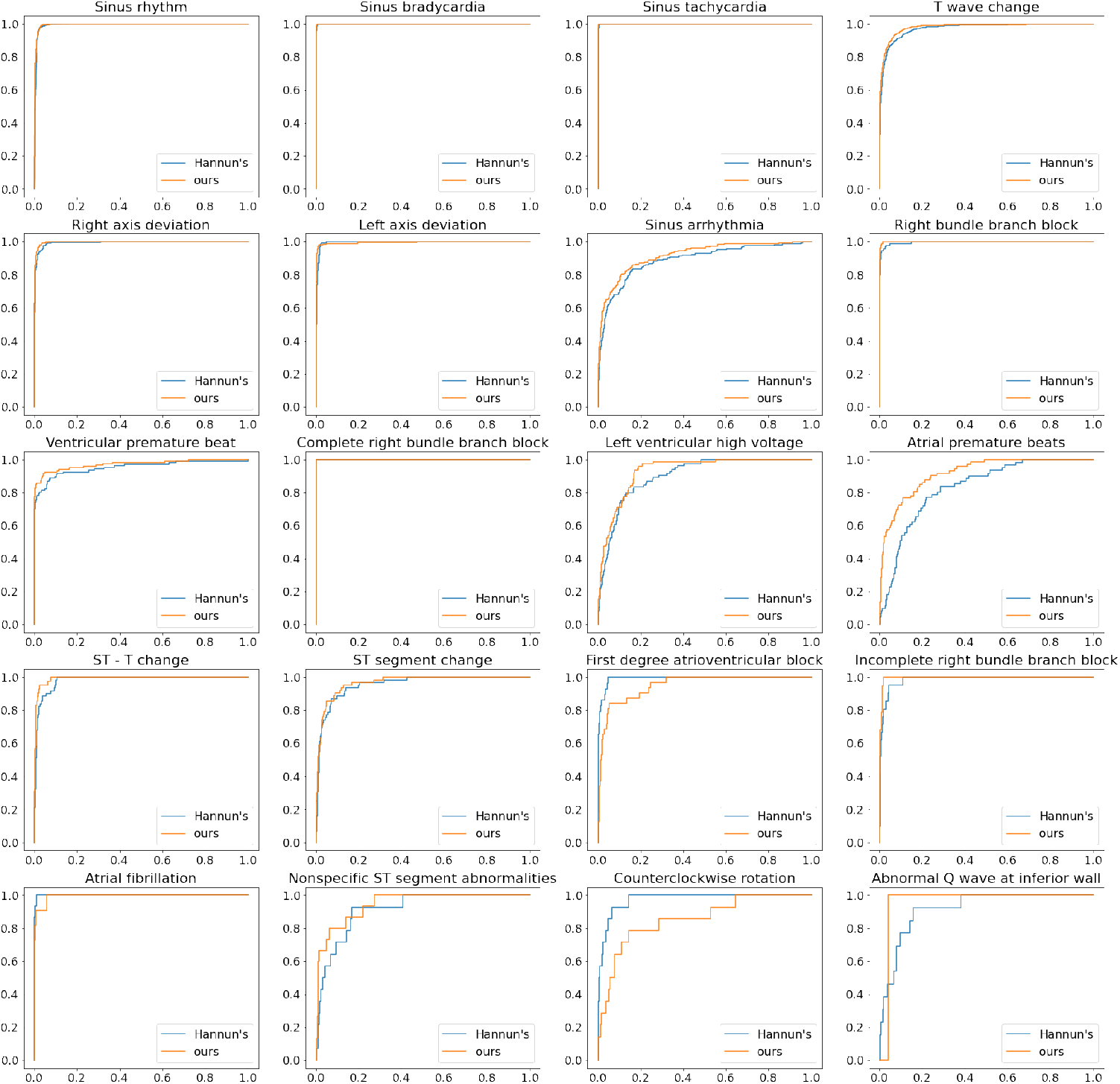
Illustrating the ROC curves of our proposed method and the Hannun’s [9]. Notably, our proposed method outperform the Hannun’s [9] in the most types of disease diagnosis.

### D. Ablation Study

To inspect the contributions of different modules, we conduct an ablation study, as presented in Table II. The proposed modules are all beneficial to ECG signal diagnosis. Notably, versions with more or fewer Encoder modules (reported in Line (3)–(6)) cannot obtain the performance gains, which might be because non-local features are relatively comprehensively learned. Besides, our method can achieve comparable performances even without one particular module. With more Encoder modules, prediction precision increases, but recall decreases. This phenomenon may be attributed to an overfitting issue where too many Encoder modules overfit to tiny features.

**TABLE II.** The ablation study of the proposed modules.

| Methods | Precision | Recall | Micro F1 |
| --- | --- | --- | --- |
| Ours (original) | 0.943 | 0.890 | 0.916 |
| (1) w/o SE-Attention | 0.939 | 0.889 | 0.913 |
| (2) w/o Temporal Attention | 0.938 | 0.888 | 0.912 |
| (3) w/ 1 * Encoder module | 0.940 | 0.888 | 0.913 |
| (4) w/ 3 * Encoder module | 0.943 | 0.889 | 0.915 |
| (5) w/ 4 * Encoder module | 0.943 | 0.885 | 0.913 |
| (6) w/ 5 * Encoder module | 0.947 | 0.877 | 0.911 |

### E. Visualizations

To evaluate whether the proposed classifier extracts the essential features, we have adapted the GradCAM method [27] for the ECG signals and visualized the critical segments in the automatic diagnosis, as depicted in Fig. 5. The proposed method focuses on specific segments, whereas ResNet34 highlights the entire cardiac cycle. This visualization reinforces the notion that our method effectively captures essential features.

**Fig. 5.**
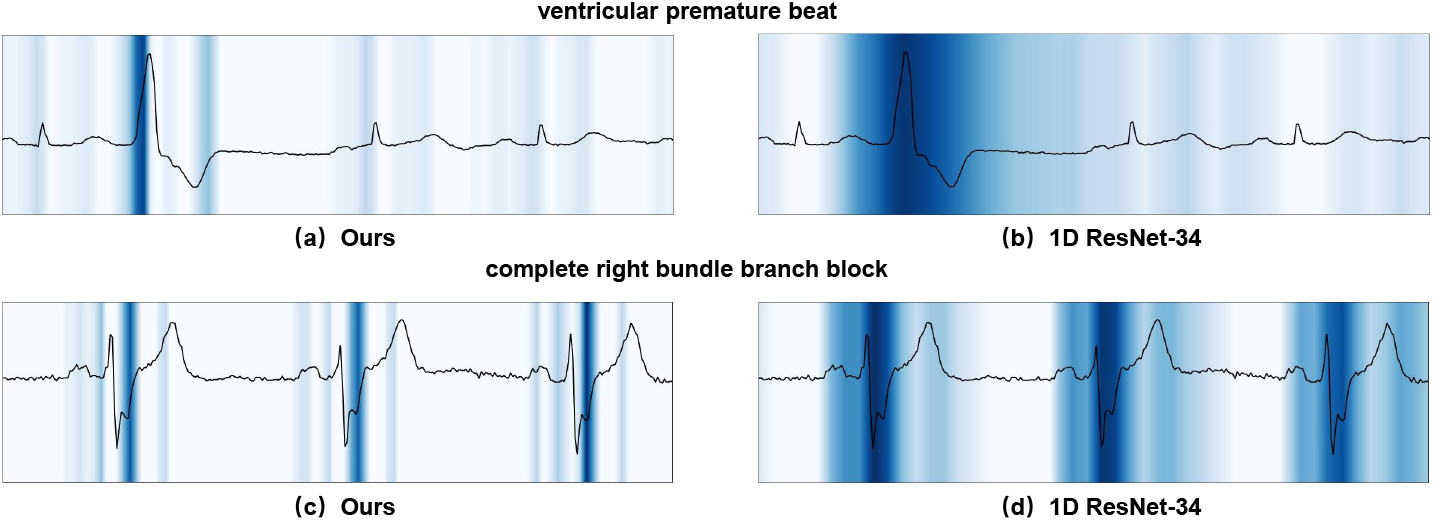
Visualization Comparisons between our approach with convolutional neural networks (using 1D ResNet as example).

## V. Conclusions

In summary, this paper has introduced a neural classifier utilizing self-attention operations to model long-dependencies in time signals for ECG signal diagnosis. Our approach initially fuses features from various leads using the Lead Fusing block. Subsequently, we employ Transformer-XL encoders with sliding windows to capture non-local features and further learn global features. The final step involves a fully connected layer for ECG signal diagnosis (classification). Experiments conducted on the Tianchi ECG dataset have demonstrated that our method outperforms state-of-the-art approaches. Ablation study provides insights into the effect of each module, and GradCAM visualizations underscore the interpretability of our approach while maintaining strong performance.

## Acknowledgment

The study is supported by the Startup Fund for Scientific Research, Fujian Medical University (Grant number: 2022QH1268).

## Footnotes

1 https://ecglibrary.com/norm.php

2 https://tianchi.aliyun.com/competition/entrance/231754/information?lang=en-us

3 https://tianchi.aliyun.com/competition/entrance/231754/rankingList/1

